# Genomics of primary chemoresistance and remission induction failure in pediatric and adult acute myeloid leukemia

**DOI:** 10.1101/051177

**Authors:** Fiona C. Brown, Paolo Cifani, Esther Drill, Jie He, Eric Still, Shan Zhong, Sohail Balasubramanian, Dean Pavlick, Bahar Yilmazel, Kristina M. Knapp, Todd A. Alonzo, Soheil Meshinchi, Richard M. Stone, Steven M. Kornblau, Guido Marcucci, Alan S. Gamis, John C. Byrd, Mithat Gonen, Ross L. Levine, Alex Kentsis

## Abstract

Despite intense efforts, the cure rates of children and adults with AML remain unsatisfactory in large part due to resistance to chemotherapy. Whilst cytogenetic risk stratification proved valuable in identifying causes of therapy failure and disease relapse, cytogenetically normal AML remains the most prevalent disease type, with significant heterogeneity of clinical outcomes, including primary chemoresistance. Using targeted sequencing of 670 genes recurrently mutated in hematologic malignancies, we investigated the genetic basis of primary chemotherapy resistance and remission induction failure of 107 primary cases obtained at diagnosis from children and adults with cytogenetically normal AML. Comparative analysis revealed mutations of *SETBP1, ASXL1* and *RELN* to be significantly enriched at diagnosis in primary induction failure as compared to remission cases. In addition, this analysis revealed novel genomic alterations not previously described in AML, as well as distinct genes that are significantly overexpressed in therapy resistant AML. However, identified gene mutations were sufficient to explain only a minority of cases of primary induction failure. Thus, additional genetic or molecular mechanisms must cause primary chemoresistance in pediatric and adult acute myeloid leukemias.

**Key Points:**

- Targeted gene sequencing of 670 genes in adult and pediatric AML
- Profiling of 107 primary AML samples identifies new genomic alterations for primary chemoresistance

## Introduction

For patients with acute myeloid leukemia (AML), failure to achieve complete remission after induction therapy or relapse after complete remission represents the major barrier to cure for both children and adults. Despite recent efforts to improve risk stratification and to incorporate targeted therapies into AML care, relapse rates remain approximately 30% and 50% for children and adults with AML, respectively^1–3^. Whilst leukemia cytogenetics at the time of diagnosis has proven to be the most useful prognostic biomarker of clinical outcomes and therapy stratification, patients with cytogenetically normal (CN) AML encompass the largest group for whom prognostication and therapy selection are hindered by the lack of prognostic markers and effective therapeutic targets.

Genomic profiling has been used to stratify patients with CN AML, identifying recurrent somatic gene mutations, such as those in *NPM1*, *TET2*, *ASXL1*, and *DNMT3A*, that are significantly associated with clinical outcomes^4^, ^5^. Recent studies using whole-genome sequencing have assessed the mutational landscape of relapsed AML in adults identifying two major patterns of clonal evolution in response to chemotherapy: 1) persistence of the dominant pre-leukemic clone with mutations acquired at relapse, and 2) expansion of a diagnostic sub-clone, emphasizing the importance of therapeutically targeting specific molecular mechanisms that contribute to chemotherapy resistance^6–8^. Similar observations were recently made using genomic studies of pediatric AML, with specific variant allele frequencies representing potential prognostic indicators^9^. Clonal persistence after induction chemotherapy appears to be associated with the increased risk of relapse^10^, but whether these mutant clones directly cause chemoresistance or reflect age-related clonal hematopoiesis remains to be determined^11^. Importantly, the molecular basis of primary chemotherapy resistance, and thus fundamental causes of chemoresistance in general, remain poorly understood. Here, we used targeted gene sequencing of the majority of genes known to be recurrently mutated in hematologic malignancies to identify diagnostic gene mutations that are associated with primary chemoresistance in adult and pediatric CN AML.

## Study design

### Patient Selection

Specimens were collected from patients treated at the Memorial Sloan Kettering Cancer Center, MD Anderson Cancer Center, Dana-Farber Cancer Institute, Ohio State University Comprehensive Cancer Center, and Children’s Oncology Group (COG) participating institutions. All patients provided informed consent, were enrolled on respective institutional research protocols and treated using current institutional remission induction regimens (Table 1). Primary chemoresistance was defined based on the presence of at least 5% AML blasts by morphologic assessment of bone marrow aspirates obtained after two cycles of induction chemotherapy, as assessed by the respective institutional or central pathologic review.

**Table 1:** Demographic features of induction failure and induction remission cohorts

| Characteristics | Pediatric |  | Adult |  |
| --- | --- | --- | --- | --- |
|  | Remission | Failure | Remission | Failure |
| Number, N (%) | 25 (48) | 27 (52) | 32 (58) | 23 (42) |
| Gender, % female | 52 | 44 | 47 | 39 |
| Age (years), median (range) | 11.5 (2.8-16.8) | 10.4 (0.6-16.8) | 48.4 (19-84) | 54.7 (16-73) |
| Therapy, N (%) |  |  |  |  |
| ADE | 25 (48) | 27 (52) | 0 | 0 |
| IDA + HDAC | 0 | 0 | 10 (31.2) | 4 (17.4) |
| 7+3 DNR and AraC | 0 | 0 | 11 (34.4) | 14 (60.9) |
| FAI | 0 | 0 | 4 (12.5) | 1 (4.3) |
| BID - FA | 0 | 0 | 2 (6.25) | 2 (8.7) |
| FLAG - IDA | 0 | 0 | 4 (12.5) | 0 (0) |
| IDA + AraC | 0 | 0 | 1 (3.1) | 1 (4.3) |
| Clofa + IDA + AraC | 0 | 0 | 0 | 1 (4.3) |

Abbreviations: ADE, cytarabine, daunomycin and etoposide-based regimen; IDA, idarubicin; HDAC, high-dose cytarabine; DNR, daunorubicin; BID-FA, twice daily fludarabine + cytarabine; FAI, fludarabine, cytarabine, idarubicin; FLAG-IDA, fludarabine, cytarabine, idarubicin and granulocyte colony stimulating factor; Clofa, clofarabine.

### Targeted Gene Sequencing

Mononuclear cells isolated from bone marrow aspirates obtained at diagnosis were purified by Ficoll gradient centrifugation, and enriched for leukemia cells by negative immunomagnetic selection against CD3, CD14, CD19 and CD235a given the lack of expression of these markers by the majority of AML cells^12^. Targeted DNA and RNA capture and sequencing of 670 genes (405 genes in DNA and 265 genes in RNA) known to be recurrently mutated in hematologic malignancies were performed using the Foundation One Heme sequencing platform. Extraction of DNA and RNA, library preparation and Illumina sequencing were carried out as previously described^13^. Sequencing data were analyzed to identify genomic alterations including base substitutions, insertions, deletions, and gene fusions^13^. Differential gene expression analysis was performed based on per million processing regions (PPM) gene expression measurements, as median normalized to all other AML specimens analyzed to date^13^. All statistical comparisons were corrected for multiple hypothesis testing using the Bonferroni correction, and statistical significance was calculated as indicated.

### Results and Discussion

We were able to obtain genomic profiles of 107 CN AML specimens in total, collected from 55 adults and 52 children, balanced for outcome groups, gender and age (Table 1). No significant differences were observed in the total number of coding mutations between the primary failure of induction chemotherapy and complete remission outcome groups for both pediatric and adult cohorts (Supplemental Table 1). Consistent with previous studies, the most frequently mutated genes among adults were *DNMT3A* (49%), *FLT3* (42%) and *NPM1* (42%), and for pediatric patients *FLT3* (54%), *NRAS* (27%), and *WT1* (25%), which were not significantly associated with either outcome group (Figure 1). Mutations of *DNMT3A* and *NPM1* were relatively rare in pediatric (2%) as compared to adult cases (*p* < 0.001, permutation test). In contrast, *CEBPA* mutations were more frequently observed in pediatric as compared to adult CN AML (21% versus 2%, respectively, *p* < 0.001, permutation test).

**Figure 1:**
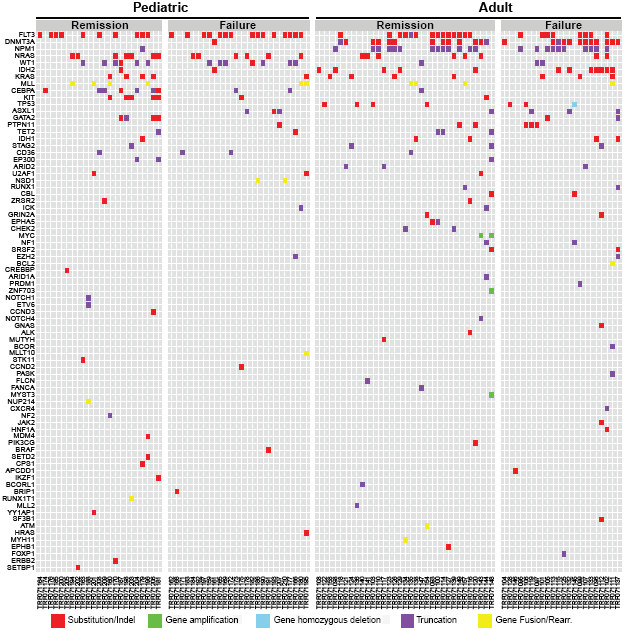
Mutational spectrum of pediatric and adult AML with primary chemotherapy resistance and induction failure. Tile plot of the genomic alterations identified in DNA/RNA targeted sequencing. Data are shown for 107 CN AML specimens, collected from 52 and 57 adult and pediatric specimens respectively, grouped by clinical outcome.

Genomic profiling also identified 15 gene rearrangements among the 21 specimens for which diagnostic material of adequate quality was available for RNA capture (Supplemental Table 1). Cryptic rearrangements of known genes in AML were detected, including *KMT2A (MLL)* and *NUP98-NSD1*. The latter was observed in only pediatric patients from this study, consistent with previous reports^14^, ^15^. We identified several novel cryptic gene rearrangements including intergenic deletions involving *NOTCH1/EDF1* and *GATA2/DNAJB8*, as well as translocations leading to the rearrangements of *SQSTM1-NUP214* and *IGH-BCL2*. Mutations of these genes have been reported in hematologic malignancies, for example the *SQSTM1-NUP214* gene fusion in cases of T-cell acute lymphoblastic leukemias (ALL), but not in AML^16^. In addition, our study identified potential novel coding mutations in *MAP3K1* (*MEKK*), *MAP3K6* (*MEKK6*), *EP300* (*p300*), and *WDR90* (Supplemental Table 1). These findings emphasize the value of genomic profiling to identify potential pathogenic lesions that may elude conventional cytogenetic analyses and diagnosis based on morphologic or immunophenotypic studies.

We assessed whether any gene mutations were significantly associated with primary chemoresistance as compared to complete remission among the adult and pediatric patients in the studied cohorts. This identified mutations of *SETBP1* (8 out of 50 versus 1 out of 57, *p* = 0.01) and *ASXL1* (7 out of 50 versus 1 out of 57, *p* = 0.02), as well as *RELN* (8 out of 50 versus 2 out of 57, *p* = 0.04, permutation test). Mutations of *SETBP1* and *ASXL1* have been reported to be associated with inferior outcomes and increased risk of relapse in patients with AML^17^, ^18^. Relapse-associated *SETBP1* mutations in AML commonly cause missense mutations in the SKI domain (p.858-871). Our analysis identified the missense p.D868N mutation in one CN AML specimen, as well as additional missense mutations (p.E183K, p.A374T, p.A408T, p.M502I, p.D1161H, p.T1547N and p.S1590C) of SETBP1, suggesting that alternative mechanisms of SETBP1 dysregulation may exist in AML. Mutations of *RELN* have been reported in early T-cell precursor (ETP) ALL, a stem cell-like leukemia with high rates of intrinsic chemotherapy resistance and genetic features that are shared with subtypes of AML^19^. Our findings of *RELN* mutations associated with primary chemoresistance in CN AML raise the possibility that these leukemias may comprise a distinct biologic entity.

In addition, using available RNA sequencing and gene expression data, we assessed whether genes included in our analysis were significantly altered in expression in the induction failure as compared to complete remission cohorts (Supplemental Table 2). This analysis revealed significant overexpression of genes encoding the DNA damage repair and response regulators *ATM*, *ATR*, *PSIP1 (LEDGF)*, and *EMSY (C11ORF30)*, endoplasmic reticulum stress response regulator *HERPUD1*, transcription factors *CBFB* and *MEF2C*, kinases *AKT3* and *MAP3K7 (TAK1)*, metabolic enzymes *LDHA* and *ACSL6*, clathrin *CLTC*, and immune regulator *CTLA4* (*p* < 0.05, t-test). Additional profiling and functional studies will be necessary to establish their prevalence and pathogenicity in AML. Indeed, recent studies have implicated overexpression of *MEF2C* with inferior outcomes in pediatric AML^20^, and overexpression of *MAP3K7* has been observed to enhance AML cell survival^21^. These results suggest that other mechanisms such as those involving epigenetic or functional molecular alterations of cell signaling may also mediate chemotherapy response and failure.

In summary, we have assessed the mutational landscape of cytogenetically normal AML at diagnosis in adults and children to determine the genetic basis of induction failure and primary chemoresistance. In spite of careful selection of cytogenetically normal specimens, our study identified numerous cryptic gene rearrangements, including those that have potential therapeutic implications. Likewise, we found novel gene alterations including lesions that have been observed in non-myeloid hematologic malignancies, some of which may be pathogenic. In addition, our analysis revealed significant overexpression of genes that are associated with primary chemoresistance. Though mutations of *SETBP1*, *ASXL1* and *RELN* appear to be associated with primary chemoresistance and induction failure in our study, their prevalence was relatively low. We did not identify single gene mutations that are significantly associated with primary chemoresistance and induction failure in the majority of the patients studied. It is possible that additional genetic lesions such as those not included in the current target gene panel^13^, or alternatively, combinations of genetic alterations that are below the statistical power of our study may cause primary chemoresistance and failure in AML, which may be elucidated by larger and more comprehensive genomic profiling studies. Finally, it remains to be determined whether functional molecular profiling such as that achieved using gene expression, epigenetic and proteomic analyses may reveal more universal biomarkers and therapeutic targets to prospectively identify and therapeutically block chemotherapy resistance in AML^22–25^.

## Acknowledgements

We thank Alejandro Gutierrez for critical discussions. This work was supported by the NIH R21 CA188881, K08 CA160660, P30 CA008748, U10 CA180899, Burroughs Wellcome Fund, Alex’s Lemonade Stand Foundation, Gabrielle’s Angel Foundation, and the Josie Robertson Investigator Program.

## Authorship

FB and AK wrote the manuscript and performed analysis. FB, PC, ES performed laboratory experiments. JH, SZ, SB, DP and BY performed analysis. KK, TA, SM, RS, SK, GM, JB compiled patient samples. ED, MG and JH performed statistical analysis. AK and RL designed the study. RL contributed clinical and genomic expertise.

## Conflict-of-interest disclosure

JH, SZ, SB, DP and BY are employees and equity holders of Foundation Medicine Inc. RL is a consultant for Foundation Medicine Inc.

## Correspondence

Alex Kentsis, MD, PhD, Memorial Sloan Kettering Cancer Center, 1275 York Ave, New York, NY, USA 10065, Phone number: 1-646-888-2593

---

**Supplemental Table 1:Genomic alterations**.

**Supplemental Table 2:Gene expression analysis**.

